# Plant-pollinator interaction linkage rules are altered by agricultural intensification

**DOI:** 10.1101/2020.01.10.900878

**Authors:** Beth M. L. Morrison, Berry J. Brosi, Rodolfo Dirzo

## Abstract

Determining linkage rules that govern the formation of species interactions is a critical goal of ecologists, especially considering that biodiversity, species interactions, and the ecosystem processes they maintain are changing at rapid rate worldwide. Species traits and abundance play a role in determining plant-pollinator interactions, but we illustrate here that linkage rules of plant-pollinator interactions change with disturbance context, switching from predominantly trait-based linkage rules in undisturbed, natural habitats, to abundance-based linkage rules in intensive agricultural habitats. The transition from trait-based to abundance-based linkage rules corresponds with a decline in floral trait diversity and an increase in opportunistic interaction behavior as agricultural intensification increases. These findings suggest that agricultural intensification is changing the very rules determining the realization of interactions and the formation of communities, making it challenging to use the structure of undisturbed systems to predict interactions within disturbed communities.

## Introduction

Biotic interactions play a crucial role in determining the ecological and evolutionary trajectories of species and the ecosystem processes they maintain (Bailey et al., 2012; Berlow et al., 2009; Guimarães, Pires, Jordano, Bascompte, & Thompson, 2017; Naeem, Duffy, & Zavaleta, 2012; Pilosof, Porter, Pascual, & Kéfi, 2017; R. Thompson et al., 2012; Tylianakis, Laliberté, Nielsen, & Bascompte, 2010; Vázquez & Aizen, 2004). In a world of increasing human development and rapid environmental change, understanding the drivers behind the formation of pairwise species interactions is a nontrivial but critical challenge for ecologists (Bartomeus et al., 2016; Berlow et al., 2009; Gray et al., 2014; Harvey, Gounand, Ward, & Altermatt, 2017; Hutchinson et al., 2019; Olito & Fox, 2015; Pomeranz, Thompson, Poisot, & Harding, 2019; R. Thompson et al., 2012). The ability to predict species interactions would enhance our understanding of the biological and functional outcomes of environmental disturbance (Gray et al., 2014; Hutchinson et al., 2019) and could consequently inform conservation schemes targeted at mitigating, preventing, and reversing the negative impacts of human disturbance on biodiversity and ecosystem processes (Harvey et al., 2017; Hutchinson et al., 2019; Kaiser-Bunbury & Blüthgen, 2015; Tylianakis et al., 2010). Furthermore, as our desire to understand the role and importance of species interactions grows, so too does our need for empirical interaction data. However, collecting interaction data is costly and time-consuming. Therefore, accurate predictive models of species interactions would be a boon to interaction network research (Bartomeus et al., 2016; Gonzales et al., 2018; Gray et al., 2014; Jordano, 2016; Kaiser-Bunbury & Blüthgen, 2015; Monteiro & Faria, 2018; Pomeranz et al., 2019). Despite the potential applications, a robust and predictive understanding of the rules that determine the formation of species interactions remains elusive.

Given that a pair of species co-occur in space and time, the realization of an interaction between them may be determined in large part by the respective traits of each species (i.e. niche processes or biological constraints; Jordano, Bascompte, & Olesen, 2003; Poisot, Stouffer, & Gravel, 2015; Santamaría & Rodríguez-Gironés, 2007). For example, whether a pollinator’s tongue can reach the nectar at the base of a flower corolla or if the gape size of a predator is large enough to eat a prey species. Indeed, some studies have found that for a suite of interaction types only a few traits are needed to accurately predict a significant number of the interactions in an interaction network (Bartomeus, 2013; Bartomeus et al., 2016; Berlow et al., 2009; Eklöf et al., 2013; Monteiro & Faria, 2018; Pomeranz et al., 2019; Santamaría & Rodríguez-Gironés, 2007). On the other hand, species abundance may also determine the linkage rules of interactions (i.e. neutral processes; Canard et al., 2012; Poisot et al., 2015; Vázquez & Aizen, 2004), as the relative abundance of species has been found to accurately predict interaction network structure (Bartomeus et al., 2016; Olito & Fox, 2015; Pomeranz et al., 2019; Vázquez, Chacoff, & Cagnolo, 2009).

Prior studies that have attempted to disentangle whether traits or abundance drive interaction linkage rules have furthered our understanding of the rules defining network assembly. However, a further consideration is the ability of species to change their interaction behavior, and hence the linkage rules themselves, based on community composition or resource availability. Particularly in disturbed areas, where species richness and trait diversity are typically reduced (Clavel, Julliard, & Devictor, 2011; Flynn et al., 2009; Forrest, Thorp, Kremen, & Williams, 2015), generalist species predominate (Aizen, Morales, & Morales, 2008; Aizen, Sabatino, & Tylianakis, 2012; Devictor et al., 2008; Gámez-Virués et al., 2015; Grass, Berens, Peter, & Farwig, 2013; Layer, Hildrew, & Woodward, 2013; Le Viol et al., 2012; Marrero, Torretta, Vázquez, Hodara, & Medan, 2017; Munday, 2004; Weiner, Werner, Linsenmair, & Blüthgen, 2014) and some species may even alter their interaction behavior to interact opportunistically with abundant species. Conversely, in undisturbed, natural habitats, trait diversity is retained and co-evolutionary history remains intact, which we predict could result in predominantly trait-based interaction behavior, though no studies have addressed this directly. Whether disturbance context changes the influence of traits or abundance on linkage rules remains largely unexplored, yet it will be a critical determinant of whether linkage rules can be used to predict interaction formation in novel and disturbed habitats.

Here, we use plant-pollinator interaction networks from 16 sites along a gradient of agricultural intensification (Morrison, Brosi, & Dirzo, 2019) to explore whether linkage rules of plant-pollinator interactions change with disturbance context. We chose to focus on a gradient of agricultural intensification as it is the dominant driver of global change (IPBES, 2019) yet it has considerable potential for supporting biodiversity and ecosystem services (Kremen & Merenlender, 2018). First, we assess if linkage rules are predominantly determined by plant traits or floral abundance at each site using a hierarchical modeling approach (Bartomeus, 2013). The hierarchical model estimates the probability of detection of each plant-pollinator interaction and allows for the incorporation of species traits or abundance as co-variates (Bartomeus, 2013). Then, we calculate what proportion of interactions we can correctly predict for each site using abundance values alone. Finally, we assess whether there are corresponding declines in plant trait diversity and increases in opportunistic pollinator interaction behavior as agricultural intensification increases.

## Materials and methods

The methods for data collection were originally published in Morrison et al. (2019) but we briefly summarize them here. The study took place in Santa Cruz and Monterey counties, California, U.S.A. We selected 16, 4 ha sites that fell along a gradient of agricultural intensification. These included natural sites with no history of agricultural development, organic diversified farms that featured mature hedgerows of native plants and multiple crop types, and industrial, organic monoculture strawberry farms with no diversifying features and disturbed habitat along the field margins consisting of predominantly exotic, weedy vegetation. Each site received an agricultural intensification rating based on the quantity and quality of non-crop habitat within the site. Using this system, diversified sites that had a large area of hedgerow habitat (high-quality, non-crop habitat) would receive a low agricultural intensification rating, and monoculture sites with a small area of non-crop habitat that was mostly disturbed (low-quality) would receive a high agricultural intensification rating. All the natural sites received a rating of 1, the lowest possible rating. The highest rating was 3.20 and the mean ± S.D. was 1.97 ± 0.86 (see SI for further details on the equations to calculate the ratings).

We visited each site in an approximate two-week rotation between May and August (12 sites were sampled in 2016 and 4 were sampled in 2017 but there was no significant effect of sampling year on community composition, see SI). During each sampling visit, we placed 4-7, 25m x 2m transects throughout the site within areas of maximum floral diversity. The exact number of transects sampled during each visited was dependent on time and weather, as transects were only sampled in clear skies, between 10am-2:30pm, when the windspeed was below 7 m/s on average and temperatures were above 65°F. Over the full sampling season, 32 transects were sampled at each site. To sample for plant-pollinator interactions, a researcher would walk slowly back-and-forth along the length of each transect for 15 total “active” sampling minutes, stopping the timer to catch and collect any pollinator (a bee, or Syrphid or Bombyliid fly) that unambiguously came into contact with the reproductive parts of a flower.

All pollinator samples were identified to the lowest possible taxonomic unit with the help of taxonomists (Jaime Palewek and Robbin Thorpe). We identified 35 species, 25 genus, and 34 morphospecies (grouped to genus first). For analysis we treated all levels as “species”. We pooled interaction observations to create one weighted (quantitative) plant-pollinator interaction network per site.

To quantify the floral abundance at each site, we counted the number of flowers on every species within a 1m^2^ quadrat placed every 5m along the transects and pooled the data for each site. Each flower on an individual was counted, rather than counting inflorescences, as the presence and shape of inflorescences was used as a co-variate in our model. Proportional abundance values were used in all analyses. To account for plant traits, we compiled a database of 7 traits (flower color, inflorescence/solitary flower, inflorescence shape, flower shape, flower symmetry, median flower width, and family as a proxy for a conglomerate of plant traits) for each flowering plant species we observed. Plant traits were gathered from botanical descriptions in The Jepson Manual: Vascular Plants of California and Calflora.org. We selected this suite of traits as they may influence pollination interactions. See the values for trait categories in Table 1.

**Table 1.** The floral trait categories used in the hierarchical model to determine pollinator interaction linkage rules. A list of all the trait values for the seven trait categories that we input into the hierarchical species occupancy model to determine which pollinator species had trait-based interaction preferences or abundance-based preferences for interaction partners.

| Trait Category | Values |
| --- | --- |
| Family | Asteraceae, Brassicaceae, Polygonaceae, Papaveraceae, Apiaceae, Rhamnaceae, Malvaceae, Solanaceae, Rosaceae, Fabaceae, Caryophyllaceae, Convolvulaceae, Dipsacaceae, Ericaceae, Phrymaceae, Polemoniaceae, Lamiaceae, Onagraceae, Myrsinaceae, Ranunculaceae, Cornaceae, Grossulariaceae, Cactaceae, Scrophulariaceae, Plantaginaceae, Boraginaceae, Azoaceae, Cucurbitaceae, Zygophyllaceae, Oxalidaceae, Linaceae, Liliaceae, Themidaceae, Caprifoliaceae, Cistaceae, Orobanchaceae, Garaniaceae, Carophyllaceae, Portulaceae, Lythraceae |
| Main Flower Color | white, yellow, pink, orange, red, lilac, purple |
| Inflorescence Shape | panicle, raceme, compound umbel, solitary, head, axillary, cyme, spike, calyx, umbel |
| Basic Inflorescence Shape | inflorescence, solitary |
| Flower Shape | composite, rotate, papilionaceous, funnelform, tubular, urceolate, other |
| Flower Symmetry | radial, bisymmetric, bilateral |
| Median Flower Width | continuous: 0.8 - 150 mm |

To assess whether linkage rules were determined by traits or abundance for each pollinator species at each site, we used a hierarchical modeling approach adapted from Bartomeus (2013). The model is a species occupancy model (function ‘pcount’ from the package unmarked version 0.12.2) that uses observations to predict the probability of detection of a species within a given habitat (Fiske & Chandler, 2011). Habitat attributes can be added to the model as co-variates. In our application of the model, as in Bartomeus (2013), the “habitat” is a flowering plant species and the “attributes” are the plant traits and floral abundance. The model then uses the observation information (frequency counts of each interaction) and co-variates to evaluate the best predictors of the presence of (i.e. interactions) of a pollinator. The 32 transect samples from each site were used to model the probability of detection of a pollinator within a site. Due to data limitations, we did not use these models to serve as accurate predictive models of the interactions themselves, but rather to explore whether there was evidence of an overall change in the importance of traits versus abundance in determining pollinator interactions along the gradient. A more detailed description of the application of these models to plant-pollinator linkage rules can be found in Bartomeus (2013) and a complete explanation of the model and the unmarked package can be found in Fiske and Candler (2011). All analyses were conducted in R version 3.4.2. We chose to focus exclusively on pollinator interaction partner selection rather than incorporating plant partner selection as well because pollinators can actively choose their partners, while plants have only passive partner selection. In addition, the pollinators can move across and throughout the landscape (given physiological constraints) while plants in agricultural landscapes are specifically managed and cultivated.

For each site, we created one model set for each pollinator species that was observed more than twice, resulting in 199 model sets overall. The model set included each plant trait alone as a co-variate, floral abundance as a co-variate, and all pairwise combinations of two traits. Floral trait diversity was very limited at some sites thus precluding the combination of more than two traits and the combination of traits and abundance into the same model. We then used the ‘fitList’ and ‘modSel’ functions to identify the best fit model based on AIC for each pollinator species at each site. We calculated the proportion of pollinator species that had plant traits retained as co-variates in the best fit model and the proportion that had abundance retained as a co-variate in the best fit model. We assessed the change in the proportion of species with trait-based versus abundance-based models along the agricultural intensification gradient using logistic regressions.

We reinforced our model findings and tested whether abundance values could better predict interaction identities in agricultural rather than natural areas by creating 1000 abundance-based null networks per site and calculating a change in the average number of correctly predicted interactions in each network along the gradient. To create the abundance-based networks, we used each species’ total number of interactions (from the quantitative networks) as a proxy for its relative abundance. For plants, relative abundance and number of interactions were significantly correlated (*P* < 0.0001, ρ = 0.59) and we did not have independent measures of abundance for pollinators. We created the abundance-based networks using the Patefield algorithm (function ‘nullmodel’ and method *r2dtable* in package bipartite version 2.8) which randomly shuffles the interactions from the observed network but maintains the total number of interactions per species (and thus the “relative abundance” of each species). This way, two species had a greater probability of interacting in the abundance-based networks if they had many interaction partners (i.e. were more abundant), and species with fewer interactions had a greater probability of interacting with a generalist species (representing “opportunism”). We used a logistic regression to test whether the proportion of correctly predicted present interactions in each randomized network increased with agricultural intensification. We consider only correctly predicted present interactions instead of correctly predicted absent interactions (zeros in the interaction matrix) because it is not possible to distinguish true ecologically absent interactions from interactions that were simply not observed during sampling.

There is a higher probability of correctly predicting present interactions in a network with higher connectance (the number of realized interactions in the network divided by the number of possible interactions in the network, i.e. the number of filled cells in an interaction matrix). In our study, network connectance did increase with agricultural intensification (Fig. S2). Therefore, we compared how many present interactions were correctly predicted using the abundance-based nulls versus how many were predicted in completely random nulls (50% probability of an interaction) on average. If increasing predictive accuracy of the abundance-based nulls is due to increasing connectance alone, there should be no difference between the abundance-based nulls and the completely random nulls. However, if the difference between the number of correctly predicted present interactions in the abundance-based nulls versus the completely random nulls gets increasingly larger along the gradient, this suggests that greater network connectance is not the sole driver behind the increasingly better predictive accuracy of the abundance-based nulls.

A switch to abundance-based linkage rules with increasing agricultural intensification may also be indicated by a decline in interaction specificity of pollinator species within the networks (i.e. a move from specialized interactions to generalist, opportunistic interactions). An overall decline in pollinator interaction specificity could be driven by two factors: (1) within-species decline in specificity of species found in multiple sites and habitats, and/or (2) a turnover of species along the gradient in which species with specific partner preferences (i.e. specialized species) are lost and replaced by generalist, opportunistic species.

To assess pollinator specificity, we calculated two metrics for each pollinator species at each site: (1) *d*′, a quantitative measure of specificity or reciprocal specialization (Blüthgen, Menzel, & Blüthgen, 2006), and (2) proportional degree, a qualitative measure of specialization representing the number of interactions partners each species interacted with divided by the number of possible partners within a site. Because these metrics can be affected by size and shape of the network, we standardized the values by calculating the *z*-scores using the equation 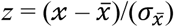 where 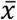 and 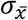 are, respectively, the mean and standard deviation of the null distribution for metric *x*. We then used a generalized linear mixed effects model with species as the random effect and agricultural intensification rating as the fixed effect to look for declines in overall pollinator specificity along the gradient. The residuals of the *d*′ *z*-scores displayed heteroscedasticity so we transformed the values by adding a constant (three) and taking the log, after which the data no longer displayed heteroscedasticity.

To look explicitly for within-species changes in interaction specificity, we also performed a mixed model on a subset of the pollinator species that occurred in at least two sites with different agricultural intensification ratings (i.e. not two natural sites; 48 species overall). Again we observed heteroscedasticity within the model residuals and we performed the same data transformation which removed the heteroscedasticity. To examine whether there was a loss of specialized species and an increase in generalist species in more intensive agricultural sites, we used a Kruskal-Wallis to compare the mean *d*′ *z*-scores of pollinator species found exclusively in natural, diversified, or monoculture sites.

Finally, a loss of plant trait diversity may be a potential driver behind any observed switch to abundance-based linkage rules in intensive agricultural habitats. Declines in plant diversity will result in fewer opportunities for pollinator species to pick their interaction partners based on preferred traits.

Therefore, we used the plant traits that we employed in the hierarchical model and explored whether plant trait diversity declined along the gradient using a random intercept mixed effects model with trait category as a random effect, agricultural intensification as the fixed effect, and the number of trait levels represented in the site in each category as the response variable.

## Results

Overall, we collected 7,945 plant-pollinator interactions, representing 95 pollinator species, 56 of which were observed at more than one site, and 141 plant species. The results of the hierarchical model revealed that the proportion of best-fit models that retained plant traits as co-variates declined along the gradient (*R*^*2*^_*adj*_ = 0.38, *P* = 0.002, *β* = -0.56, Fig. 1A). The number of correctly predicted present interactions in the abundance-based null networks increased along the gradient (*R*^*2*^_*adj*_ = 0.57, *P* < 0.001, *β* = 0.40, Fig. 1B). In addition, the mean difference between the number of correct interactions predicted by the abundance-based versus the completely random null networks got significantly larger with agricultural intensification (*R*^*2*^_*adj*_ = 0.56, *P* < 0.001, *β* = 0.02, Fig S2). This suggests that the increasing number of correctly predicted present interactions by the abundance-based nulls was not exclusively due to increasing network connectance along the gradient. Furthermore, a greater number of correctly predicted present interactions in agricultural sites is evidence that the increase in abundance-based linkage rules identified by the hierarchical models was not simply due to a decline in floral trait representation within the models.

**Figure 1.**
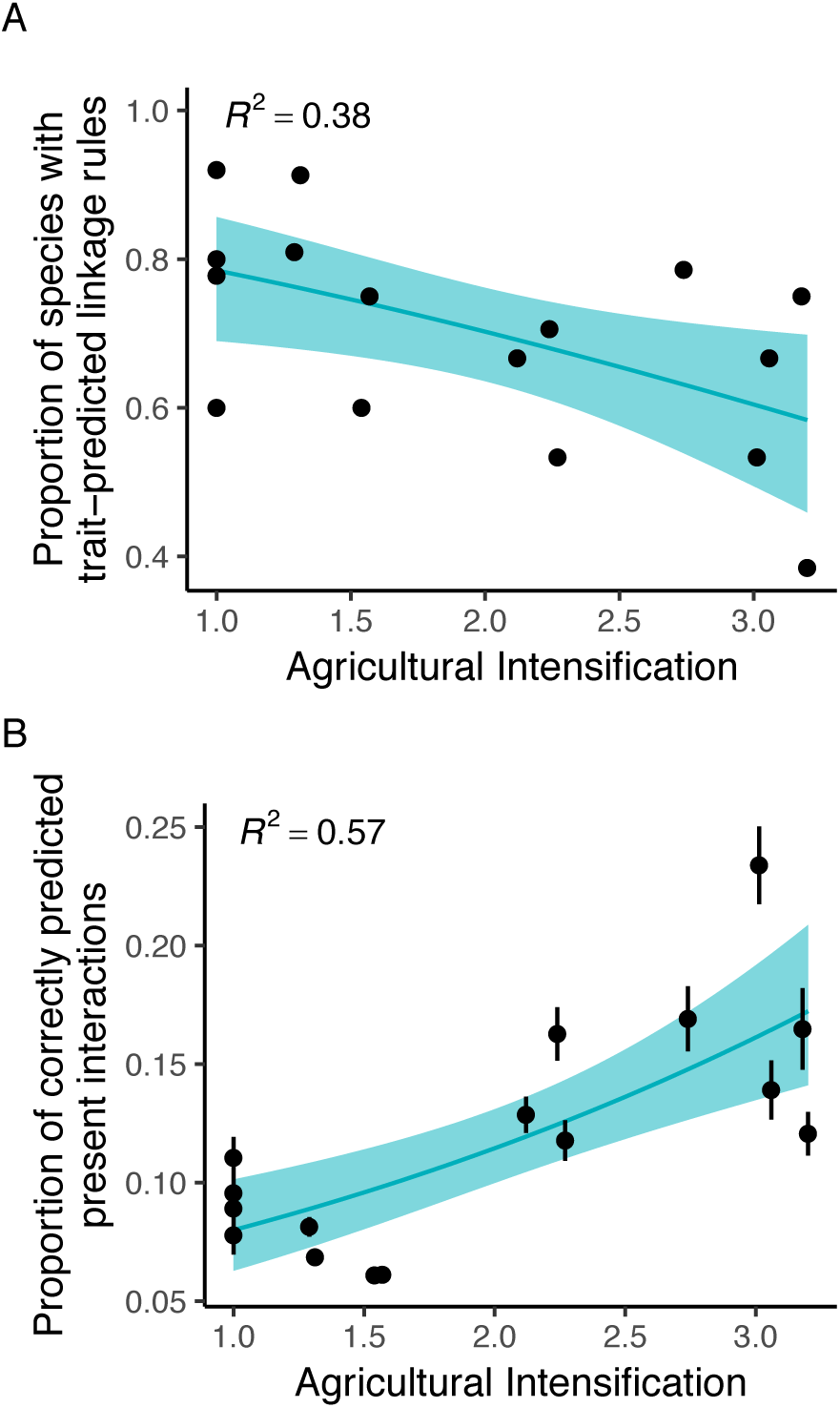
Linkage rules of plant-pollinator interaction networks changed from predominantly trait-based in natural, undisturbed areas to predominantly abundance-based in intensive agricultural areas. (A) The proportion of pollinator genera with trait-based linkage rules within each network declined as agricultural intensification increased (*P* = 0.002). Due to an increase in pollinator genera with abundance-based linkage rules, we were able to correctly predict significantly more plant-pollinator interactions in networks from more agriculturally intensive sites (B) using abundance-based null models (*P* < 0.001). Blue ribbons represent 95% confidence intervals and error bars in panel (B) represent standard deviations for 1000 null models.

We also observed a decrease in selectivity of pollinator species and an increase in opportunistic interaction behavior along the agricultural intensification gradient. The pollinator *d*′ *z*-scores declined as agricultural intensification increased (*R*^*2*^_*adj*_ = 0.32, *P* < 0.0001, *β* = -0.16, Fig. 2). Likewise, for the model that included the subset of pollinators present in at least two sites, there was a significant decline in *d*′ *z*-scores (*R*^*2*^_*adj*_ = 0.34, *P* < 0.0001, *β* = -0.17). Conversely, the proportional degree *z*-scores increased with agricultural intensification (*R*^*2*^_*adj*_ = 0.44, *P* < 0.0001, *β* = 0.60, Fig. 2) and the same trend was also observed for the subset of species present within at least two sites (*R*^*2*^_*adj*_ = 0.42, *P* < 0.0001, *β* = 0.61), demonstrating that pollinator species interacted with a greater proportion of available floral resources in agricultural sites.

**Figure 2.**
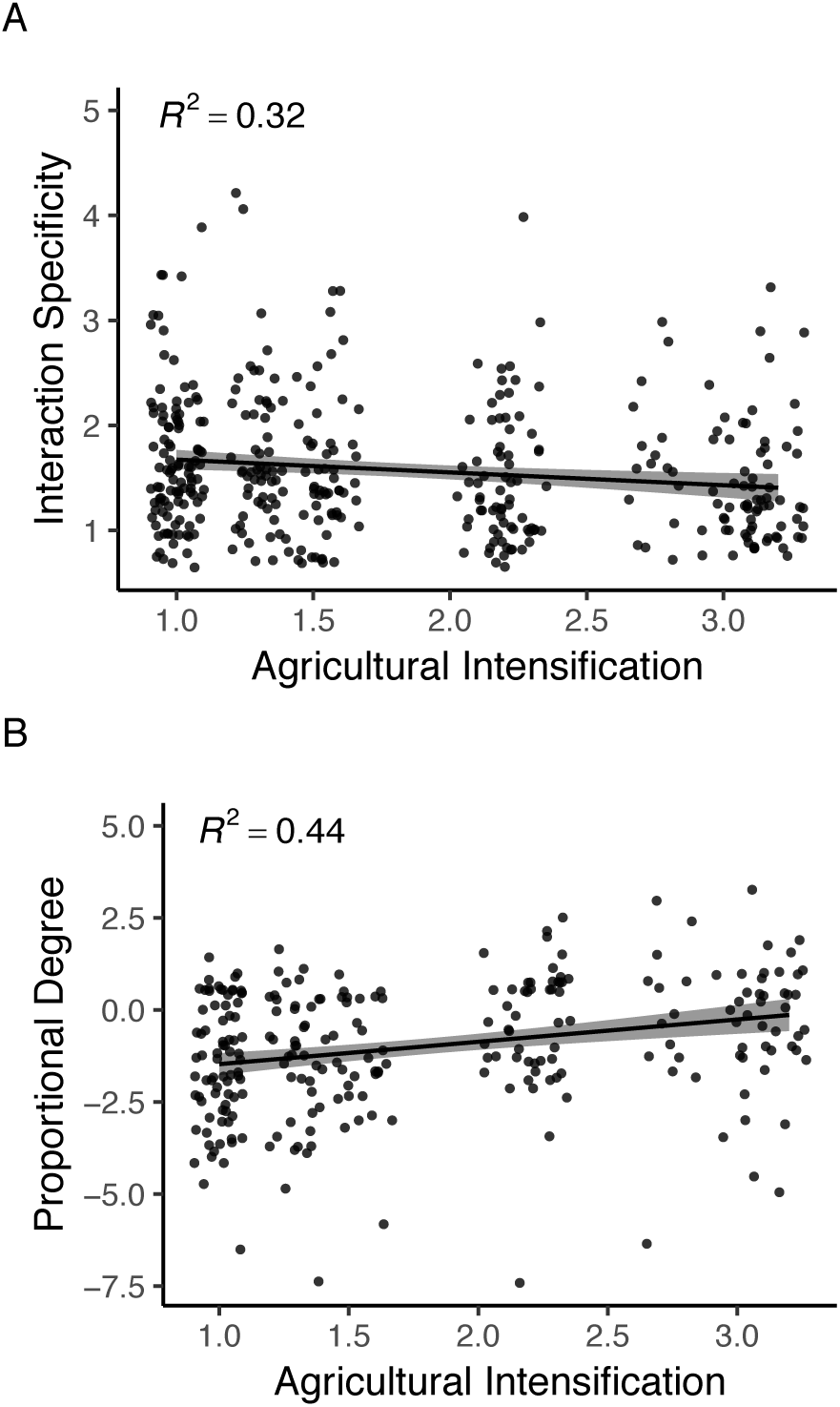
Pollinator interaction behavior became more opportunistic and less specialized as agricultural intensification increased. (A) Pollinator interaction partner specificity, measured as *d*′ *z*-scores, declined as agricultural intensification increased (*P* < 0.001), indicated that pollinators transitioned from more specialized interaction partner selection to more abundance-based opportunistic interaction partner selection. (B) The proportion of available floral resources that each pollinator species interacted with at each site also increased along the gradient (measured as proportional degree *z*-scores, *P* < 0.0001), providing further evidence that pollinators in more intensive agricultural areas are less selective in their interaction partner selection. Black ribbons represent 95% confidence intervals and data points are jittered to reduce overlap.

We did not find a significant difference among the *d*′ *z*-score values of pollinators found exclusively in natural, diversified, or monoculture habitats, though there were fewer species found exclusively in monoculture sites (9 species) versus natural (25 species) and diversified sites (20 species). This suggests that turnover of species from specialized species to generalist species as agricultural intensification increases may contribute but likely does not play a large role in the overall increase in network generalization. Lastly, plant trait diversity declined along the gradient (*R*^*2*^_*adj*_ = 0.77, *P* < 0.001, *β* = -0.89, Fig. 3).

**Figure 3.**
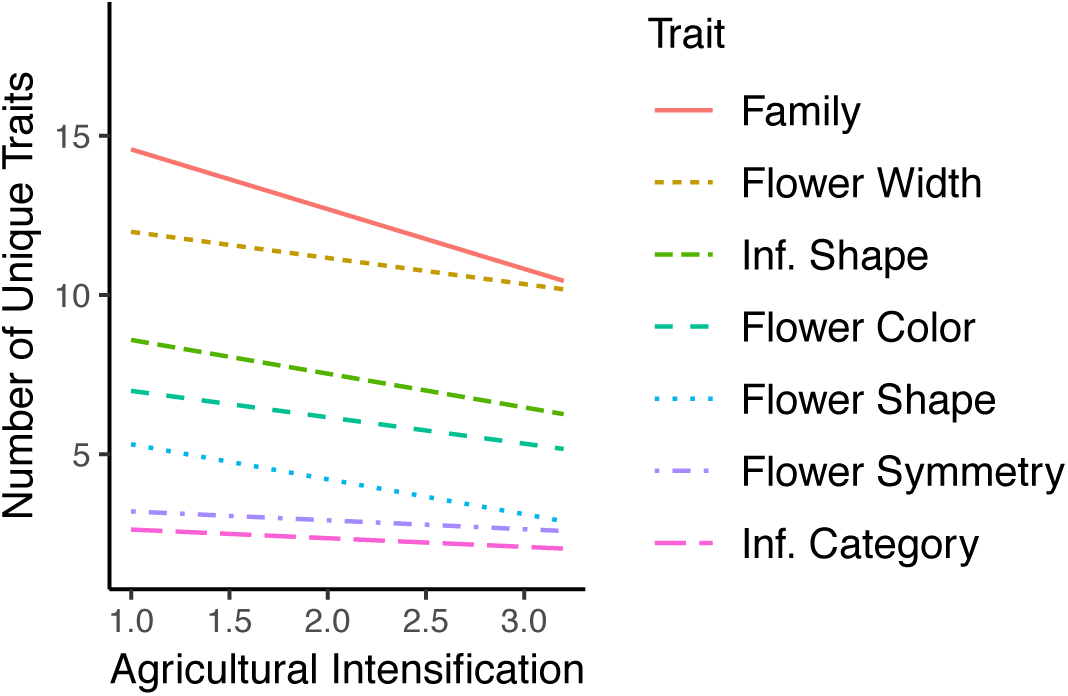
Floral trait diversity declined along the agricultural intensification gradient (*P* < 0.001). A decline in floral trait diversity may contribute to the observed increases in opportunistic interaction behavior by pollinators as they had fewer floral traits to select from in intensive agricultural areas. The data points are not shown for clarity.

## Discussion

Whether directly or indirectly, species interactions determine the assembly, persistence, and functioning of ecological communities, and understanding the processes that determine interaction realization will help ecologists predict loss and change of species interactions in the face of anthropogenic disturbance. Here we demonstrate that agricultural intensification is altering the very rules that govern the formation of species interactions within ecological communities, transitioning from primarily trait-based rules in natural habitats to abundance-based rules in intensive agricultural habitats. Our results are consistent with the hypotheses that the shift from trait-based to abundance-based linkage rules is in part due to the commensurate declines in floral trait diversity and increases in opportunistic pollinator interaction behavior along the agricultural intensification gradient.

The hierarchical model revealed that as agricultural intensification increased, the proportion of pollinator species with trait-based linkage rules declined. Accordingly, the accuracy of our interaction predictions using an abundance-based null increased along the gradient. Taken together, these results suggest that disturbance context affects the relative importance of traits versus abundance in determining plant-pollinator interaction realization. This finding furthers the conclusions of several previous studies that found that both traits (Bartomeus, 2013; Bartomeus et al., 2016; Eklöf et al., 2013; Monteiro & Faria, 2018; Pomeranz et al., 2019; Santamaría & Rodríguez-Gironés, 2007) and species abundances (Bartomeus et al., 2016; Olito & Fox, 2015; Pomeranz et al., 2019; Vázquez et al., 2009) are important for predicting the realization of species interactions. A shift towards abundance-based linkage rules in disturbed habitats was expected given that opportunistic interaction behavior is more common in disturbed habitats (Clavel et al., 2011; Devictor et al., 2008; Gámez-Virués et al., 2015; Layer et al., 2013; Le Viol et al., 2012; Munday, 2004). Because trait-based interactions often arise from the evolution of morphological or behavioral trait-matching between species or groups of species (Ehrlich & Raven, 1964; J. N. Thompson, 2005), the decline in trait-based interaction behavior in intensive agricultural habitats represents a loss of evolutionary history and has implications for the evolutionary trajectories of species in disturbed communities (Guimarães et al., 2017).

Accompanying the shifts in linkage rules were declines in pollinator species interaction specificity. Overall, the pollinators in agriculturally intensive habitats had lower interaction specificity (as indicated by the *d*′ *z*-scores) and used a greater proportion of the available floral resources than the pollinators in natural sites. If the overall decline in pollinator specificity was being driven by a loss of specialized species along the gradient, the specificity of species found exclusively in natural habitats should be higher than that of those found exclusively in agricultural habitats. Intriguingly, this was not the case, and specificity was similar between pollinators found exclusively in natural, diversified, and monoculture habitats, suggesting little species turnover form specialized to generalist species. Rather, we found that many pollinator species were present across sites and habitat types but they altered their interaction behavior to be less specific in their interaction partner choice as agricultural intensification increased. Although we did not find evidence here for the extirpation of specialized pollinators as others have (Grass et al., 2013; Weiner et al., 2014), our findings align with previous reports that pollinators are quite labile in their interaction partner selection (Brosi, Niezgoda, & Briggs, 2017; Fontaine, Collin, & Dajoz, 2008; Fontaine, Thébault, & Dajoz, 2009; Goldstein & Zych, 2016; Petanidou, Kallimanis, Tzanopoulos, Sgardelis, & Pantis, 2008; Vázquez & Aizen, 2004; Waser, Chittka, Price, Williams, & Ollerton, 1996) and that reciprocally specialized interactions are more rare in disturbed habitats (Aizen et al., 2012; Grass et al., 2013; Marrero et al., 2017). Unlike plant-pollinator interactions, other interaction types such as plant-herbivore or host-parasite interactions, are typically much more specialized between species, genera, or families (Fontaine et al., 2009; Novotny & Basset, 2005), suggesting that linkage rules for other interactions may not have the capacity to change with anthropogenic disturbance.

The decline in floral trait diversity along the gradient is a potential driver of the shift toward abundance-based linkage rules as agricultural intensification increased. Lower floral trait diversity in intensive agricultural habitats reduces the likelihood that a pollinator will find an interaction partner with its preferred floral traits. Pollinators may thus switch from interacting with their specific, preferred interaction partners to opportunistically interacting with available, but less-preferred interaction partners, resulting in a network dominated by abundance-based interactions. Other potential contributors to the shift in linkage rules could include changes in competitive behavior due to shifts in abundance and composition (Fontaine et al., 2008), changes in the nutrient landscape (requiring pollinators to broaden their diets to meet nutritional requirements) (Vaudo, Tooker, Grozinger, & Patch, 2015), or changes in “discoverability” of resources (i.e. agricultural habitats are more structurally simplified which increases encounter probability with a wider array of floral partners). Future studies that experimentally assess the potential drivers behind linkage rules would provide insight into when and why interaction realization is context dependent and its consequences for community structure.

Context-dependent linkage rules add an additional challenge to the goal of predicting the effects of anthropogenic disturbance on species interactions and the ecosystem processes they maintain. For instance, a desirable conservation goal is to *a priori* predict the effect of an environmental perturbation, such as habitat loss or the application of pesticides, on species interactions with the hopes of taking action to mitigate the negative impacts of said change (Gray et al., 2014; R. Thompson et al., 2012). We show here that the interactions in the new, post-disturbance community may not be accurately inferred based on species interaction behavior in the initial, pre-disturbance community. On the other hand, our findings indicate it may be possible to predict a proportion of the interactions within a disturbed network based on abundance values alone. This provides a promising avenue for conservation practitioners to build predicted interaction networks in disturbed or novel communities from abundance data rather than pursuing the more time-consuming collection of interaction data specifically.

Beyond disturbance context, other environmental variables likely determine the relative importance of traits and abundance on interaction realization. For example, the stability of resource availability over evolutionary time can promote the formation of reciprocally specialized interactions and the co-evolution of complementary traits between species (Dalsgaard et al., 2011; Ponisio et al., 2019; Waser et al., 1996). On the other hand, reciprocal specialization may be a less favorable strategy in unstable environments or younger communities (i.e. islands) (Ponisio et al., 2019; Waser et al., 1996), resulting in a more significant role of abundance in determining linkage rules. The evolutionary history of a community thus shapes its response to contemporary and future environmental disturbance (Rezende, Lavabre, Guimarães, Jordano, & Bascompte, 2007; Strona & Veech, 2017; Thébault & Fontaine, 2010) and determines the capacity with which interacting species are capable of altering their linkage rules to adapt to a changing environmental conditions. Furthermore, local abiotic and biotic conditions within a community, such as competition, indirect interactions, and habitat structure, also influence interaction realization (Brosi & Briggs, 2013; Poisot et al., 2015), likely with varying importance across habitat types. Overall, a strong foundation of empirical data and further exploration into the dynamic drivers behind linkage rules should inform the continued effort to build predictive models of species interactions.

## Acknowledgements

Funding for this project was provided by a Garden Club of America Centennial Pollinator Fellowship and a National Geographic Young Explorers Grant. Beth Morrison was supported by National Science Foundation Graduate Research Fellowship and an Institute for Quantitative Theory and Methods Fellowship from Emory University. Insect collections were conducted under the State of California Department of Fish and Wildlife Permit SC-13326 and specimens are stored at Stanford University. Robbin Thorpe and Jaime Pawelek assisted with pollinator specimen identification. We would like to acknowledge also the vital contributions of research assistants Samantha Faul, Hannah Nguyen, Joe Hack, Spencer Robinson, Carolynn Rice, and Samantha Baird. This project would not have been possible without the support of Amber Sciligo, David Gonthier for assistance with logistics, as well as all the farmers, land owners, and land managers who agreed to participate in this project.

