## Supplementary Information for "Plant-pollinator interaction linkage rules are altered by agricultural intensification"

### Plant-pollinator interaction linkage rules change with agricultural intensification

#### Supplementary information

##### *Study site details*

Natural sites consisted of maritime chaparral communities, a vegetation type characterized by the dominance of the woody shrubs *Arctostaphylos* spp., *Ceanothus* spp., and *Adenostoma fasciculatum*. All the diversified agricultural sites featured mature (10-15 years old) hedgerows with predominantly native species as well, including *Diplacus aurantiacus*, *Heteromeles arbutifolia*, *Eriogonum* spp., and *Ceanothus thyrsiflorus*. The monoculture and diversified sites featured many non-native, weedy species including *Brassica nigra*, *Helminthotheca echioides*, *Rumex* spp., and *Sonchus oleraceus*.

##### *Site distance*

Sites were on average 24.22 ± 14.68 km apart (min. = 2.22 km, max = 66.70 km; sites separated by minimum distance were sampled in two separate years; Fig. S1). This is a relevant distance for invertebrate pollinators. The largest and thus likely the furthest-traveling (Greenleaf, Williams, Winfree, & Kremen, 2007) pollinators we observed were *Bombus* spp. and *Xylcopa* spp. They generally travel between 1-6 km to forage (Hagen, Wikelski, & Kissling, 2011; Pasquet et al., 2008). In open landscapes of mass-flowering crops, *Bombus* can forage 11.6 km away from their colonies (Rao & Strange, 2012), but no two sites in this study were separated by completely open habitat. An ANOVA of a distance-based CCA indicated that there was no spatial autocorrelation in community composition among sites (P = 0.97, $F_{2,13} = 0.68$).

##### *Calculation of agricultural intensification*

The equations we used to determine agricultural intensification ratings of each site are thoroughly explained in Morrison et al. (2019), and we summarize them here.

We used to two equations to calculate the agricultural intensification ratings for each of the 16 sites:

$$(1) \qquad AI_{primary} = \frac{1}{\sum_{n=4}^{i} y_i \cdot \log_e x_i}$$

$$(2) \qquad AI_{final} = \frac{AI_{primary}}{\min(AI_{primary})}$$

Where $x_i$ is the percent of microhabitat $i$ within a site. The four microhabitats were defined as: natural habitat (quality ranking 4), non-crop hedgerow habitat (high quality, ranking 3), non-crop disturbed habitat (low quality, ranking 2), and crop area (lowest quality, ranking 1). The quality ranking is for each microhabitat $i$ is represented by $y_i$. $AI_{primary}$is the initial score used to calculate the $AI_{final}$ score. The natural log of $x_i$ more evenly distributes the AI ratings between the minimum and maximum agricultural intensification ratings. For this system to work, the percent of non-crop habitat must always be greater than 1%, as was the case in all of our sites. $AI_{primary}$ is calculated as the inverse of microhabitat quantity and quality so that the natural sites would receive the lowest agricultural intensification rating. In equation (2), $AI_{primary}$ is divided by the minimum $AI_{primary}$ rating so that all natural sites end up with the lowest AI rating of 1.

Originally, we planned to incorporate the proportion of natural habitat surrounding each study site into the agricultural intensification ratings, but preliminary assessments found that landscape context was not a significant predictor for observed biodiversity patterns ($P > 0.05$ for regression analyses of species richness and interaction richness against proportion of natural habitat at 100 m, 250 m, and 1 km radii).

##### *A note on farm management of the study sites*

Only three farm managers operated the seven monoculture sites in this study and all of them followed very similar protocols for pest management, including physical removal of invertebrates using bug vacuums, and periodic spraying of organic pesticides. The timing and frequency of these management techniques was dynamic, responding to need and changing throughout the season, and were beyond our control. We did not take into account the specifics of pest control management at each site because they were so similar, but also because our study was not intended to examine the effects of pest-control management on pollinator and herbivore interactions, but was rather intended to present a realistic and general picture of ecological interactions under actual management scenarios.

##### *Plant-pollinator data*

Pollinator specimens are stored at Stanford University. Males of the genus *Lasioglossum* were removed from the analysis as they are morphologically monotonous and hence unreliably identifiable (Gibbs, 2010). The *Hylaeus* genus contained at least *Hylaeus rudbeckiae* and *H. mesillae* (identified from male specimens), the females of which are also unreliably identifiable from morphological characteristics

(Snelling, 1966). Therefore, all *Hylaeus* specimens were grouped into *Hylaeus* spp. All Syrphidae and Bombyliidae were identified to genus.

##### *Floral abundance measurements*

To get measures of floral abundance, we recorded the number of flowers of each species within a $1m^2$ quadrat placed every 5m along each sampling transect. When flowers of a species were highly abundant within a quadrat, (>150) the researcher estimated floral abundance using the categories: 150-250, 250-500, 500-1000. The median of each category was used in the final measures of floral abundance. Each individual flower was counted as a floral unit, regardless of size or type of inflorescence as inflorescence shape and flower width were both included as co-variates in the model later on. The floral abundance values for each species were pooled across all quadrats from each site and the relative floral abundance of each species was used in all further analyses.

**Supplementary figures and tables**

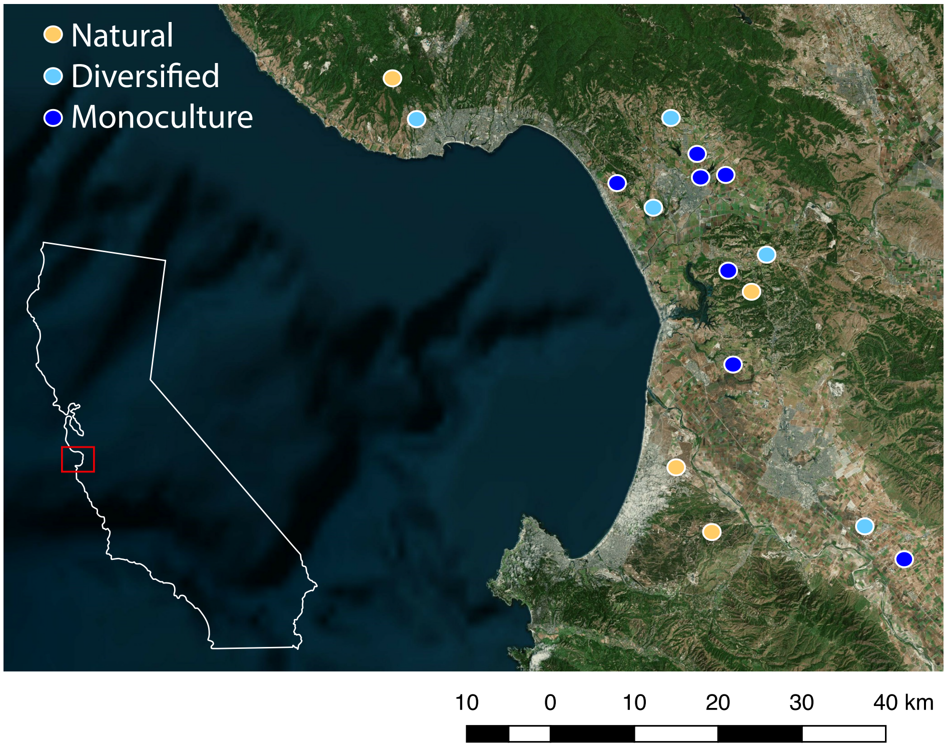


**Figure S1. The locations of the 16 study sites in Monterey and Santa Cruz counties, California, U.S.A.**

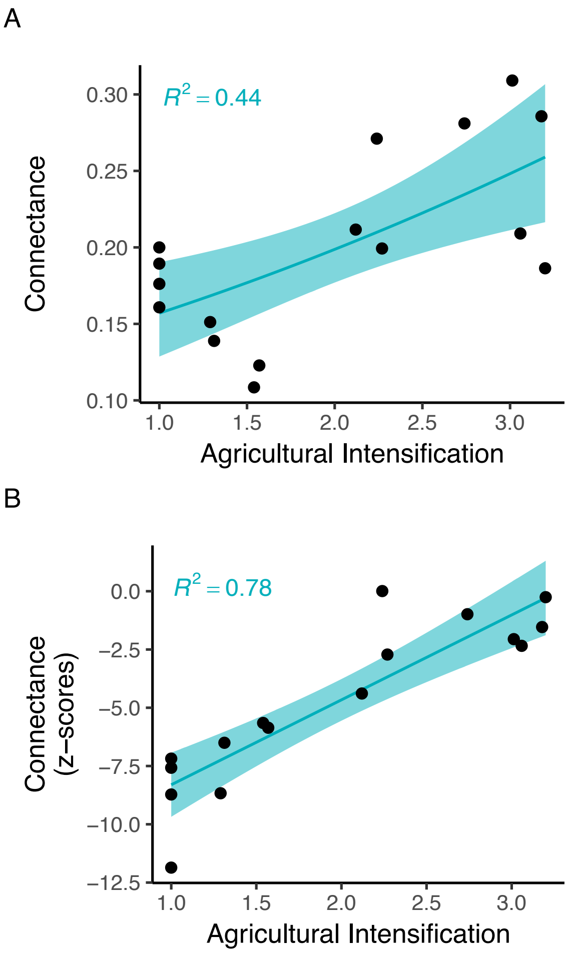


**Fig S2.** Plant-pollinator network connectance increases along the agricultural intensification gradient. The unstandardized connectance values are presented in (A) and the z-score standardized values (corrected for network size which can significantly affect connectance scores) are presented in (B). $P < 0.05$ for both regressions and blue polygons are 95% confidence intervals. Connectance *z*-scores move from significantly less connected than expected by chance in natural and less agriculturally intensive sites, to no more connected than expected by chance in agriculturally intensive sites. This further suggests that plant-pollinator interactions are more generalist, random, and less selective as agricultural intensification increases, which further supports the finding that pollinators in agriculturally intensive sites interact based on abundance (i.e. generalist, opportunistic interactions) than in natural or less agriculturally intensive sites, where they appear to be selecting partners based on traits rather than broadly and opportunistically interacting with plants in the community.

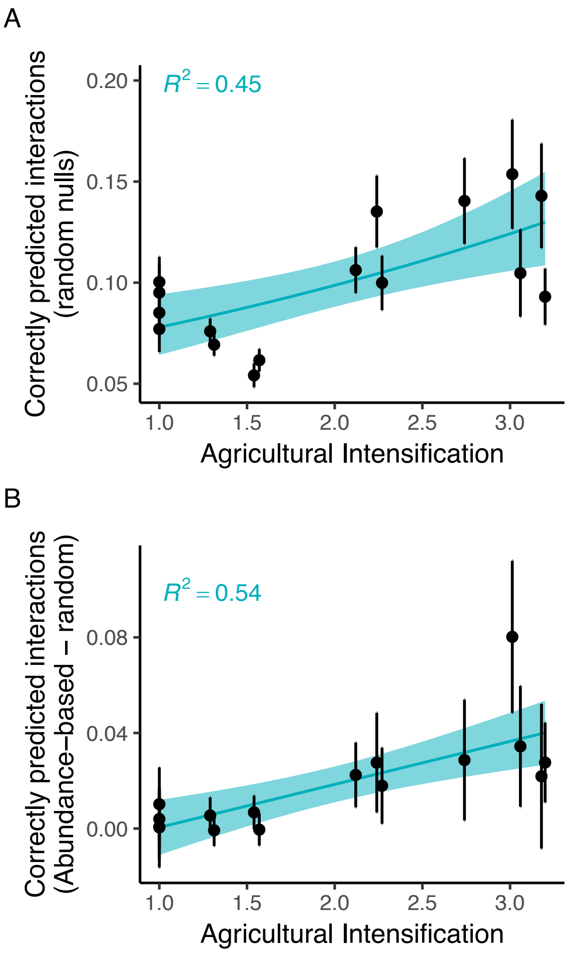


**Fig S2.** The number of correctly predicted interactions in the completely random null matrices (A). In the random matrices, each interaction had a 50% probability of being realized. The proportion of correctly predicted present interactions increases with agricultural intensification because connectance of the networks also increases with agricultural intensification (Fig. S1), and the probability of correctly predicting a present interaction increases with connectance (because more cells of the matrix are filled). However, the abundance-based null model (in which interactions were randomly distributed based on species' relative abundances in the network) increasingly outperforms the random null model at correctly predicting present interactions (B), suggesting that the increasing ability for the abundance-based nulls to correctly predict interactions is not due to increasing connectance of the networks alone. Data points are mean values from 1000 nulls and error bars are standard deviations, blue polygons are 95% confidence intervals.
